# A global barley panel revealing genomic signatures of breeding in modern cultivars

**DOI:** 10.1101/2020.03.04.976324

**Authors:** Camilla Beate Hill, Tefera Tolera Angessa, Xiao-Qi Zhang, Kefei Chen, Gaofeng Zhou, Cong Tan, Penghao Wang, Sharon Westcott, Chengdao Li

## Abstract

The future of plant cultivar improvement lies in the evaluation of genetic resources from currently available germplasm. Recent efforts in plant breeding have been aimed at developing new and improved varieties from poorly adapted crops to suit local environments. However, the impact of these breeding efforts is poorly understood. Here, we assess the contributions of both historical and recent breeding efforts to local adaptation and crop improvement in a global barley panel by analysing the distribution of genetic variants with respect to geographic region or historical breeding category. By tracing the impact breeding had on the genetic diversity of barley released in Australia, where the history of barley production is relatively young, we identify 69 candidate regions within 922 genes that were under selection pressure. We also show that modern Australian barley varieties exhibit 12% higher genetic diversity than historical cultivars. Finally, field-trialling and phenotyping for agriculturally relevant traits across a diverse range of Australian environments suggests that genomic regions under strong breeding selection and their candidate genes are closely associated with key agronomic traits. In conclusion, our combined dataset and germplasm collection provide a rich source of genetic diversity that can be applied to understanding and improving environmental adaptation and enhanced yields.

**Author summary:** Today’s gene pool of crop genetic diversity has been shaped during domestication and more recently by breeding. Genetic diversity is vital for crop species to be able to adapt to changing environments. There is concern that recent breeding efforts have eroded the genetic diversity of many domesticated crops including barley. The present study assembled a global panel of barley genotypes with a focus on historical and modern Australian varieties.

Genome-wide data was used to detect genes that are thought to have been under selection during crop breeding in Australian barley. The results demonstrate that despite being more extensively bred, modern Australian barley varieties exhibit higher genetic diversity than historical cultivars, countering the common perception that intensive breeding leads to genetic erosion of adaptive diversity in modern cultivars. In addition, some loci (particularly those related to phenology) were subject to selection during the introduction of other barley varieties to Australia – these genes might continue to be important targets in breeding efforts in the face of changing climatic conditions.

## Introduction

The diversity of the existing genetic pool for commercially important plant species has been shaped during plant domestication, human migration, varietal selection processes and, more recently, breeding. However, there is concern that breeding efforts have eroded genetic variation, thereby resulting in a narrow range of genotypes in the current gene pools of domesticated crops [1, 2]. Although regionally adapted landraces and wild relatives represent the most diverse germplasm reservoirs, the introgression of desirable alleles into elite germplasm used by breeders—whilst minimising the introduction of other genes from the wild germplasm that might reduce the agronomic fitness of the elite cultivar —has been challenging and time consuming [3]. As a result of these challenges and the often limited availability of high-density markers and detailed information for key adaptive traits, the high degree of genetic diversity in wild crop relatives has been poorly exploited.

Changes in global climate and short-term variations in growing environments pose unprecedented challenges to maintain and further enhance crop yields. Among other effects, climate change substantially alters phenological cycles, thereby posing a significant challenge to growers, who must modify crop management practices such as sowing dates in order to achieve optimal flowering times [4]. Conservation and maintenance of current crop genetic diversity for future breeding of new varieties is particularly important to help mitigate future adverse impacts of climate change on crop production.

While climate change threatens the supply of agricultural products, global demand is increasing for resource-intensive foods including meat and dairy, and alcoholic beverages including beer [5, 6]. Barley (*Hordeum vulgare* L.) is a globally important and versatile crop used for both livestock feed and brewing malts. Despite of its large and repeat-rich genome (∼5.1 Gb) distributed over seven chromosomes, it is a widely utilized diploid cereal model for genetic studies in the *Triticeae*, a botanical tribe which includes polyploid bread wheat and rye. Since the barley reference genome became publicly available [7], several genetic studies have explored the origin, domestication, and geographic spread of modern barley [8, 9], and showed that the Fertile Crescent is the main centre of domestication and genetic diversity.

Population genetic studies have examined a variety of aspects of barley genetic variation on a global scale, and have identified a striking degree of variability in traits related to flowering time, grain yield, and tolerance to abiotic and biotic factors [10–14]. This suggests that current barley germplasm resources might be harnessed to meet future challenges imposed by climate change.

Although it was only in the late 18^th^ century that barley was first introduced to Australia [15], it is currently one of the world’s largest barley producers (http://faostat.fao.org). The first introduced cultivars were poorly adapted, late-maturing, European barleys, and were susceptible to the hot, dry conditions typical of Australia. It was not until the 1960s that first Australian breeding programmes were established that specifically targeted different barley-producing regions in Western Australia, South Australia, Victoria, New South Wales, Queensland and Tasmania. Only then new breeding material from North Africa and North America was introduced to improve disease resistance against powdery mildew and cereal cyst nematodes as well as phenological adaptation of Australian barley varieties, which was further aided by the development of molecular marker technologies in the 1980s. However, the genetic impact of these breeding efforts and the extent of genetic diversity within current cultivated barley germplasm reservoirs is poorly understood.

In this study, we assessed the contributions of both historical and recent breeding efforts towards local adaptation and crop improvement in a global barley panel, using Australian barley as a model for the profound impact that breeding efforts can have on developing a previously poorly adapted crop suitable to local environments. In order to accomplish this, we analysed the distribution of genetic variants in terms of geographic region and historical breeding category and determined the genomic regions under selection and underlying candidate genes that may have been shaped by selective breeding. Associations identified between both known and novel genes and agronomic traits demonstrate the value of our global barley panel for both fundamental and applied studies. Lastly, both our selective sweep analyses and our genome-wide association studies (GWASs) highlight targets for future gene and functional allele discovery.

## Results

### Barley genomic diversity

To examine the origins and patterns of genetic diversity within the currently available barley cultivar gene pool, we assembled a global panel of 632 genotypes to represent barley genotypes from major global barley breeding programmes, including both historical cultivars and modern cultivars from 43 countries (S1 Fig, S1 File). The panel of 632 genotypes geographically diverse barley cultivars was genotyped using target capture [10, 11], low-coverage whole-genome sequencing (WGS), and genotyping-by-sequencing (GBS) by Diversity Arrays Technology (DArTseq). In total, 15,328 single-nucleotide polymorphisms (SNPs) and insertions and deletions (InDels) were detected via low-coverage WGS, 4,260 SNPs via target-enrichment sequencing, and 18,551 SNPs via DArTseq were distributed across 5,171 barley genes. The mapping of genetic markers from all 632 genotypes onto the current barley reference genome sequence [7] (IBSC v2) revealed 38,139 high-confidence genetic variants. A total of 33,486 filtered genetic markers (32,645 SNPs and 841 InDels) with a minor allele frequency (MAF) > 0.01 were used in the present study (S1 Table). As expected for target capture and DArTseq analyses (which focused on actively transcribed genes), the distribution of genetic variants across the seven chromosomes exhibited a visible gradient across chromosome compartments, from distal regions with relatively high gene density to pericentromeric regions with fewer genes (S2 and S3 Figs).

For a more detailed analysis of the genomic changes that have occurred during the history of Australian barley breeding, Australian genotypes were separated into four historical subgroups based on release date: Category A (historically relevant cultivars used in 20^th^ century breeding programmes in Australia, released between 1903 and 1998), Category B (modern cultivars with specific regional adaptations that were released between 1999 and 2005 as a result of focused barley breeding programmes across different Australian states), Category C (most recently released elite cultivars, released between 2006 and 2019); and Category D (unreleased breeding and research lines). A detailed description of the Australian varieties, including the year of release and breeder, is provided in S1 File.

The polymorphism information content (PIC) was estimated to evaluate the frequency of nucleotide variants across the entire barley population, as well as in subpopulations based on geographical regions or historical subgroups of Australian barley. From the genetic variant data for all 632 accessions in the barley panel, the PIC was estimated to be 0.17 (Table 1), although we observed marked differences among different geographical regions (Australia, Africa, Asia, Europe, North America, and South America) and among historical subgroups of Australian barley. Within historical subgroups of Australian cultivars, the observed mean PIC values were slightly higher for varieties released between 2006 and 2019 (CatC) and unreleased research and breeding lines (CatD) (both 0.16) than for varieties released between 1903 and 1998 (CatA, 0.14) or between 1999 and 2005 (CatB, 0.15).

**Table 1:** Summary of molecular diversity and polymorphism information content for the whole panel and all subgroups.

| Group | Average MAF | PIC | N |
| --- | --- | --- | --- |
| Whole panel | 0.14 | 0.17 | 632 |
| 2-row group | 0.14 | 0.17 | 579 |
| 6-row group | 0.15 | 0.18 | 48 |
| Spring | 0.14 | 0.17 | 520 |
| Winter/facultative | 0.14 | 0.17 | 41 |
| Australian (all) | 0.13 | 0.16 | 227 |
| Australian (CatA) | 0.12 | 0.14 | 16 |
| Australian (CatB) | 0.13 | 0.15 | 14 |
| Australian (CatC) | 0.13 | 0.16 | 17 |
| Australian (CatD) | 0.13 | 0.16 | 180 |
| European | 0.13 | 0.16 | 141 |
| North American | 0.14 | 0.17 | 183 |
| South American | 0.13 | 0.16 | 34 |
| Asian | 0.15 | 0.18 | 24 |
| African | 0.13 | 0.15 | 15 |

MAF: Minor allele frequency. PIC: Polymorphism information content. N: Number of genotypes per group (where information available, see S1 File). CatA: Cultivars released between 1903 and 1998, CatB: cultivars released between 1999 and 2005; CatC: cultivars released between 2006 and 2019; Cat D: breeding and research lines.

### Population structure within the global collection of domesticated barley varieties

Underlying population structure is known to be a confounding factor in GWASs, particularly for adaptive traits such as flowering time [16]. Known sources of population structure in domesticated barley varieties include the separation of two-row and six-row barleys, which occurred early in domestication (∼8,000 years ago), as well as the separation of spring and winter barleys, which accelerated the migration of barley through the modification of the vernalisation requirement and photoperiod response [17, 18]. Next, we therefore investigated the population structure of the global barley germplasm collection used in this study using ADMIXTURE [19] to select the optimal number of subpopulations (K), which we predicted was approximately K = 12 (S4 and S5 Figs; Figs 1a–d).

Phylogenetic trees were constructed based on the genetic distances of the entire population (Figs 1e and f), as well as on the genetic distances of 47 Australian cultivars selected to represent the diversity of germplasm used in Australian barley breeding (S6 Fig) using the Neighbour-joining (NJ) clustering method. Distinct clusters were detected based on geographic location (Fig 1e), row type, and growth habit (Fig 1f). As expected, no clear clustering pattern was observed based on historical subgroups (S6 Fig), as within our Australian barley panel, many of the cultivars that were first to be released are ancestors of modern cultivars. Principal component analysis (PCA) was performed with separation based on row type, growth habit, or geographic region (S7a–c Figs), corroborating the results of phylogenetic analyses. Taken together, our data suggest that three major factors account for the partitioning of diversity within the global barley panel: geographic origin (Asia, Middle East, North America, and South America) (S7a Fig), growth habit (winter vs. spring) (S7b Fig), and row type (six-row vs. two-row) (S7c Fig). However, no clear clustering pattern was observed among European, African, and Australian genotypes, which is likely due to the extensive movement of germplasm from Europe to Australia and extensive use of diverse African lines in Australian and European breeding programmes [15].

**Figure 1:**
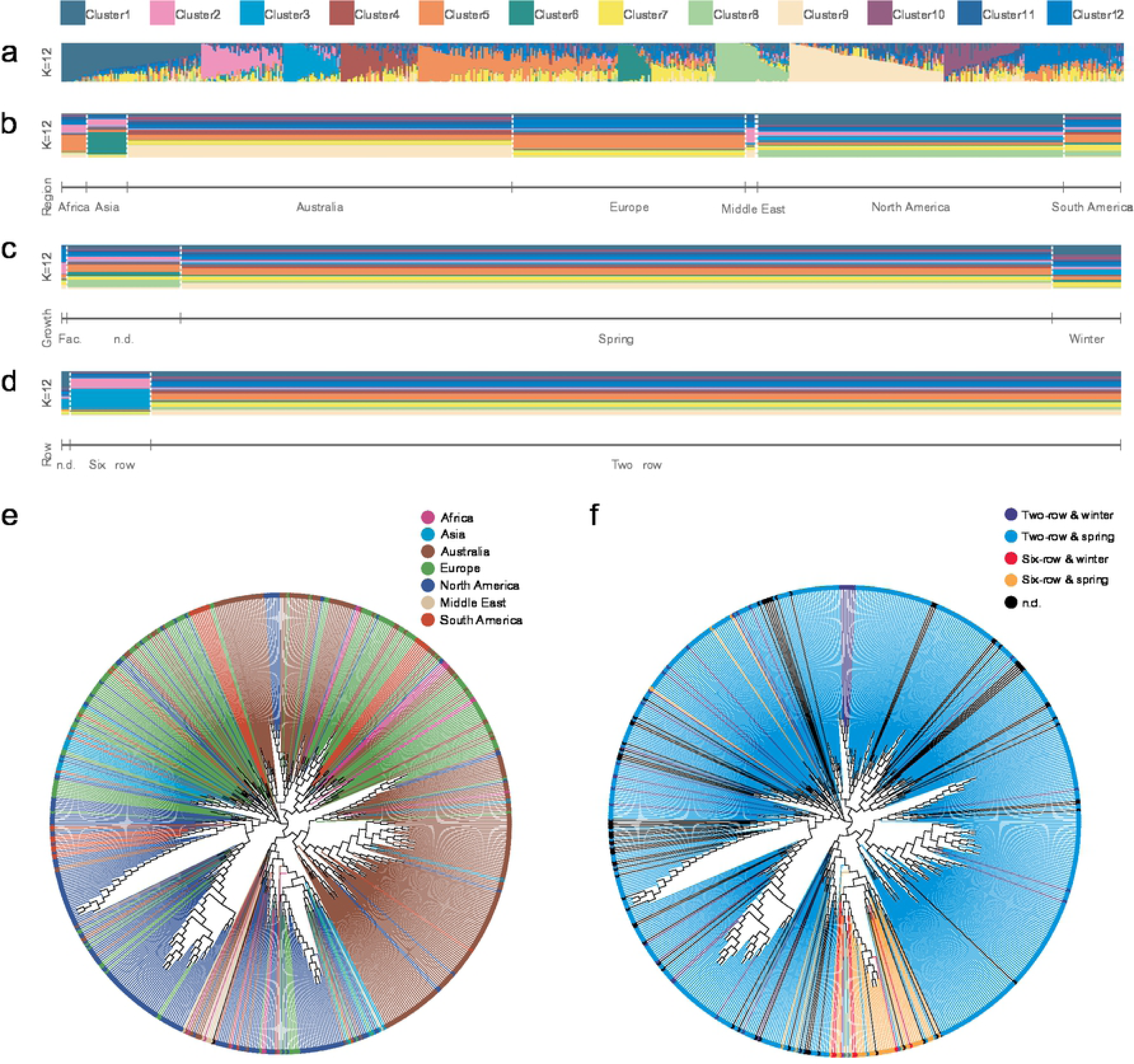
Population structure of the global barley panel. a) Population structure of the entire barley panel was inferred by assuming twelve subpopulations (K) (**Supplemental Figure 4**). Each colour represents a different subpopulation as per the legend. Distribution of ADMIXTURE-defined populations based on b) seven geographical locations, c) three growth habits, and d) two row types. The neighbour-joining trees of 632 barley genotypes with clusters highlighted are based on e) geographic location or f) growth habit. The trees were constructed from simple matching distances of 33,486 common genetic variants in the barley population. Fac., facultative.

To understand the patterns of linkage disequilibrium (LD) between different chromosomes, we calculated *r^2^* values between pairs of genetic variants for all 632 genotypes, as well as in the four Australian subpopulations (S8 Fig). LD was estimated for each subpopulation as a function of physical distance (Fig 2).

**Figure 2:**
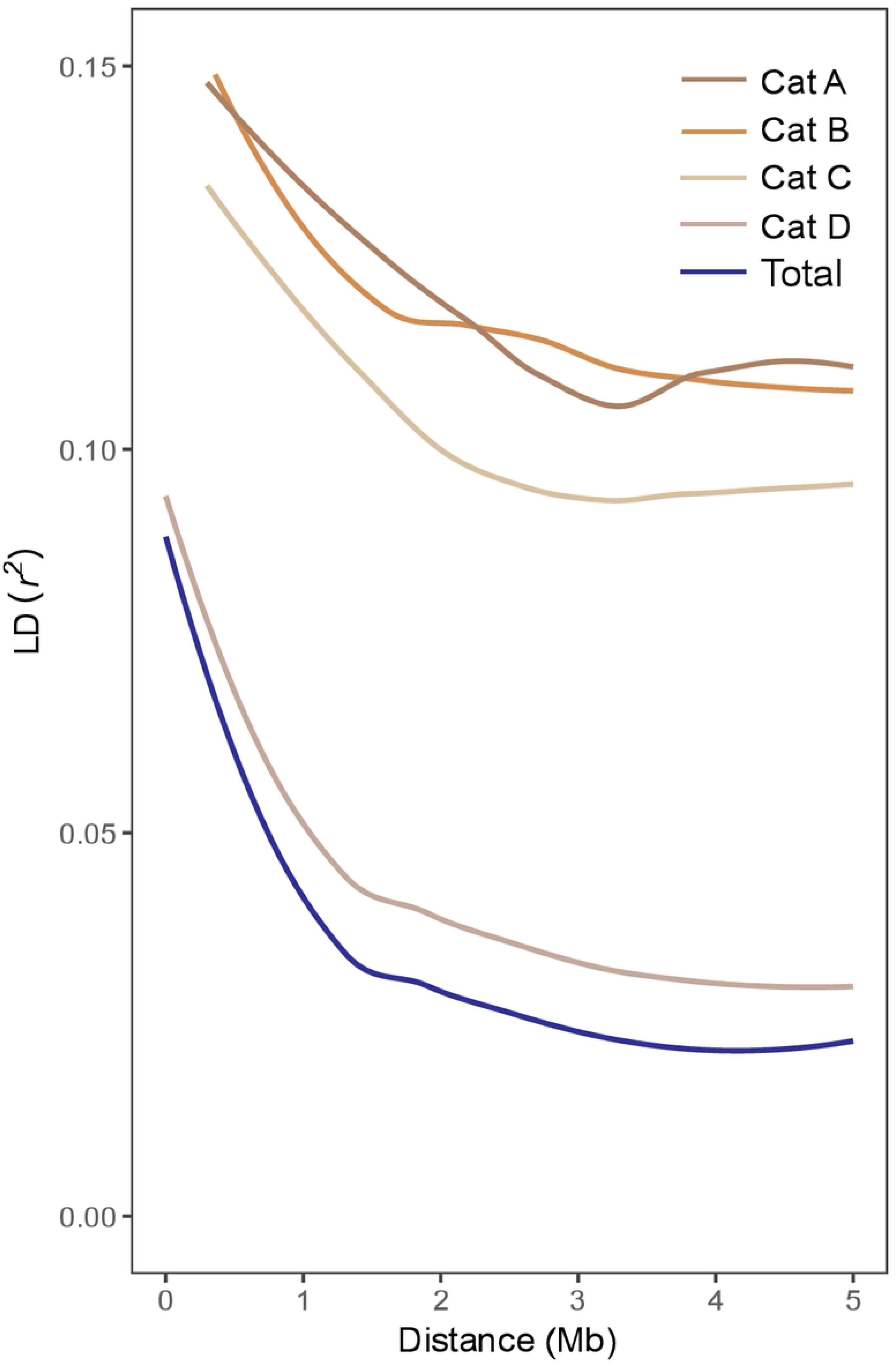
Genome-wide linkage disequilibrium (LD) decay in different historical groups of domesticated Australian barley genotypes. Values are reported as mean LD *r^2^* for all pairs of genetic variants binned by distance (100 kb). Curves were fitted by a LOESS function. CatA: Cultivars released between 1903 and 1998, CatB: cultivars released between 1999 and 2005, CatC: cultivars released between 2006 and 2019, Cat D: breeding and research lines, Total: total barley population of 632 varieties.

Genetic marker pairs were sorted into 100-kb bins based on the distance between pairs, and mean *r^2^* values were estimated for each bin (S2–6 Files). Owing to selection pressure on large genomic regions for positive alleles, the subsequent fixation of the alleles during breeding, and high rates of self-fertilization, Australian barley subgroups (CatA to CatC) were found to contain larger LD blocks, higher baseline LD, and higher long-range LD than the entire barley panel used in this study. Long-range LD was more extensive in historical barley cultivars (CatA and CatB) than in more recently released barley cultivars (CatC) owing to the greater extent of allelic association in the early period of barley breeding, thereby confirming the narrow initial gene pool of early breeding programmes [15].

### Selection footprints of barley breeding

To explore selection footprints resulting from breeding within the global barley panel, next we investigated genetic diversity parameters in for the whole population, between barley genotypes sourced from different geographic regions, and within Australian subgroups based on release date. We first examined the degree of polymorphism along each chromosome within different geographic region and among historical Australian barley groups by calculating the nucleotide diversity statistic π [20] (S9 and S10 Figs). The distribution of nucleotide diversity indicates limited allelic diversity in domesticated barley genotypes and that modern breeding processes had measurably altered overall genetic diversity, which increased during a relatively short breeding period in Australian barley varieties (Fig 3a, S2 Table).

**Figure 3:**
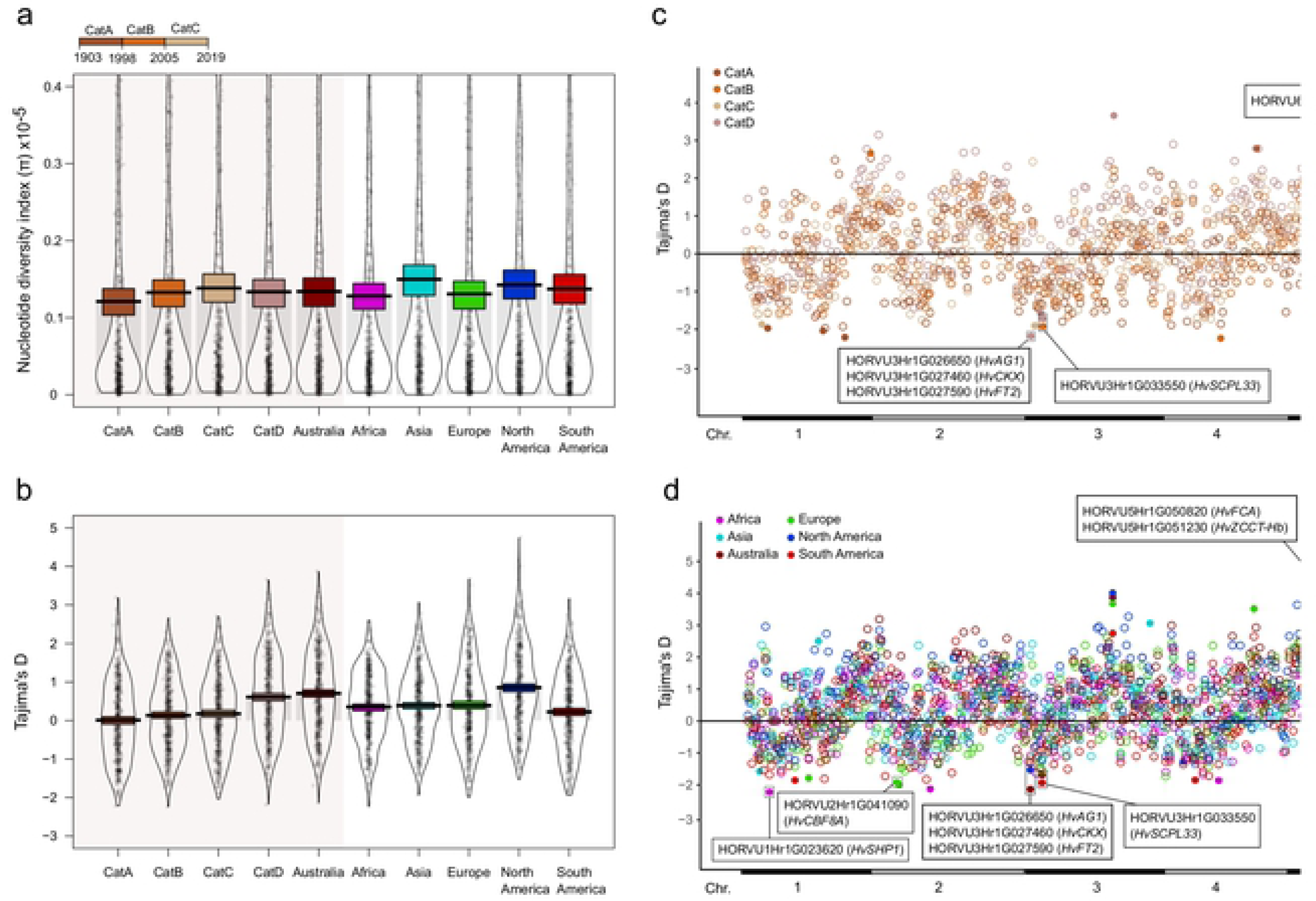
Genetic diversity and selection (breeding) signatures of different groups of domesticated barley genotypes. a) Plots of nucleotide diversity index (π) values and b) Tajima’s D values to compare the average number of pairwise differences and the number of segregating sites between samples within each of our geographic and historical subpopulations in Australia (highlighted in light grey shading; a timescale is provided above the panel). Solid thin black horizontal lines indicate means, transparent horizontal bands of different colours indicate Bayesian 95% highest-density intervals (HDIs), black dots represent individual data points, full densities are shown as bean plots. c) Tajima’s D distribution among the different historical groups of domesticated Australian barley genotypes and d) barley varieties from different geographic regions. Filled circles show values above the 99^th^ percentile and are colour coded according to the different historical or geographic groups as indicated in the legends within the panels. Boxes point to data points above the 99^th^ percentile that are located within phenology-related genes. Details are further described in the figure. All statistics are based on 10-Mb windows. CatA: Cultivars released between 1903 and 1998; CatB: cultivars released between 1999 and 2005; CatC: cultivars released between 2006 and 2019; Cat D: breeding and research lines.

The nucleotide diversity index π varied between barley varieties from the six geographic regions, with the highest and lowest genetic diversities observed in Asian and African barleys, respectively. A gradual increase in nucleotide diversity was detected when comparing historical cultivars (CatA) to the later (CatB) and the most recently released Australian cultivars (CatC). Our results are consistent with a continuous increase in diversity through breeding, as observed between CatA and CatB (representing a ∼8.9% higher nucleotide diversity in cultivars released between 1998 and 2005 than in cultivars released between 1903 and 1998). The continuous increase in diversity between groups CatA and CatC (representing a ∼12.5% higher nucleotide diversity in cultivars released between 2005 and 2019 than in cultivars released between 1903 and 1998) reflect the increased use of exotic germplasm bred into modern Australian barleys [15] and breeding improvement in early Australian barley breeding programmes. These findings show that despite six decades of intense breeding of Australian barley cultivars, higher genetic variation exists within the current breeding gene pool compared to historical varieties.

We also calculated subpopulation-specific estimates of Tajima’s D to compare the average number of pairwise differences and the number of segregating sites between samples within each of our geographic and historical subpopulations in Australia. The sign of Tajima’s D provides an interpretation of natural selection, where balancing selection results in a positive, and positive selection results in a negative Tajima’s D. Subpopulation-specific estimates of Tajima’s D differed extensively, and included both negative and positive values, but with a strong and consistent skew towards positive mean values for all subpopulations (Fig 3b, S11 and S12 Figs). In addition, we observed an excess of rare alleles relative to expectation (corresponding to negative Tajima’s D values in the top 1% tail of the empirical distribution) for five phenology-related genes in historical Australian barley groups and seven phenology-related genes in geographic subpopulations, including known phenology genes *FLOWERING LOCUS T2* (*HvFT2*) and *AGAMOUS 1* (*HvAG1*) in North American and in CatD Australian subpopulations (Figs 3c and d). We also observed positive Tajima’s D values, indicating a lack of rare alleles relative to expectation, which corresponds to a sudden population contraction, likely associated with the introduction of barley varieties to Australia and other countries (Figs 3c and d). Positive Tajima’s D values were detected for phenology-related genes in historical Australian barley groups, including *FLOWERING LOCUS T1* (*HvFT1*) for the earliest-(CatA) and latest-released (CatC) Australian barley cultivars (Fig 3c), as well as for the North American and Asian varieties (Fig 3d).

To unravel genomic regions targeted by breeders in efforts to improve barley production in Australia in the last 120 years, we next explored loci in the barley genome that harbour selective sweeps related to breeding. To accomplish this, we examined population differentiation using the fixation index (F_ST_) (S13 Fig), reduction of diversity (ROD), and cross-population composite likelihood ratio [21] (XP-CLR) test scores, and compared these results within the Australian panel between the groups CatB and CatC groups (which have been subjected to recent breeding efforts) and those for the oldest group, CatA (Fig 4, S13–15 Figs).

**Figure 4:**
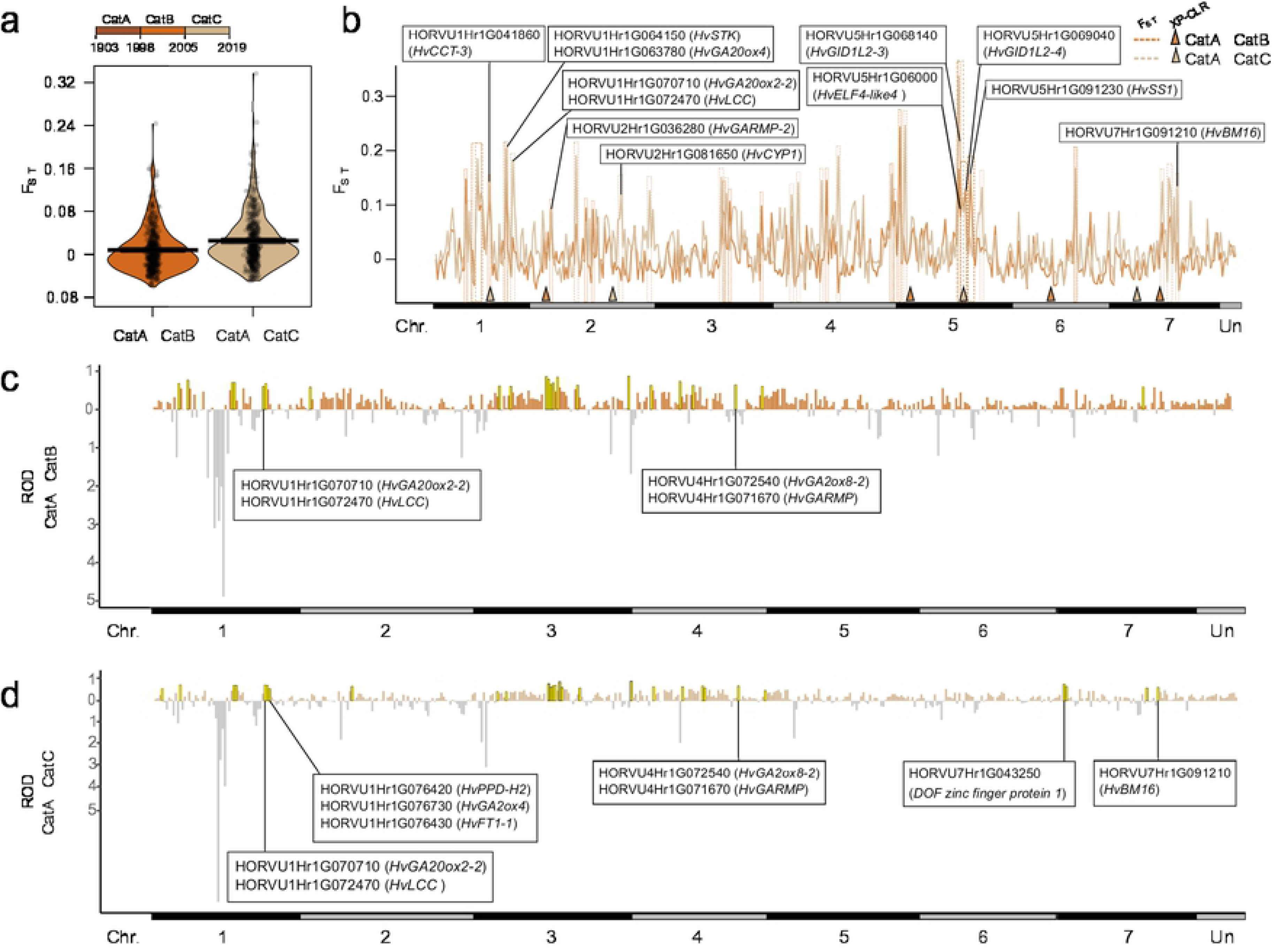
Breeding selection signatures of domesticated Australian barley genotypes. a) Pirate plot of the genetic differentiation fixation index (F_ST_) values between different historical groups of domesticated Australian barley genotypes (a timescale from 1903 to 2019 is provided above the panel). Solid thin black lines indicate means, black horizontal bands indicate Bayesian 95% highest-density intervals (HDIs), black dots represent individual data points, full densities are shown as bean plots. b) F_ST_ values and cross-population composite likelihood ratio (XP-CLR) test scores for CatA, CatB, and CatC historical barley groups on each chromosome (Chr.), illustrating the range of variation in diversity between these groups. c) Reduction of Diversity (ROD) distribution between the CatA and CatB historical barley, and d) distribution between the CatA and CatC historical barley groups. Highlighted regions (as per the legend for b and yellow bars for c and d) are above the 95^th^ percentile (F_ST_ and ROD), or above the 99^th^ percentile (XP-CLR). Boxes indicate regions located within phenology-related genes, with details further described in the figure. All statistics are based on 10-Mb windows. CatA: Cultivars released between 1903 and 1998, CatB: cultivars released between 1999 and 2005, CatC: cultivars released between 2006 and 2019.

We identified substantial population differences (high F_ST_, Figs 4a and b) and genomic regions with substantially lower levels of diversity in more recently released cultivars than in historical cultivars (CatB and CatC groups, high ROD, Figs 4c and d) as possible candidate regions that were under selection during breeding in the recent past. To further assess the extent of genetic differentiation between early and recent Australian barley cultivars, we also used likelihood ratio (XP-CLR) tests [21] to identify genomic regions that had been differentially selected between the groups (Fig 4b). Regions above the 99^th^ percentile of XP-CLR selection signals were considered candidates that had undergone selection during breeding, revealing 8 regions from the individual comparisons (CatA–CatB and CatA–CatC), of which 6 were adjacent to high-F_ST_ loci (Fig 4b). Based on the XP-CLR analysis, regions that had undergone selection contained 459 variants from 4 genes, with different regions between the two historical subpopulations.

In total, we identified 69 candidate regions with 922 potential genes that were potentially under selection during crop breeding, post-domestication and diversification in Australian barley (S7 and S8 Files). Among those genes, we identified 17 unique phenology-related genes, including gibberellin metabolism-related genes (the gibberellin oxidases *HvGA2ox8*, *HvGA20ox2*, *HvGA20ox4*, and *HvGA2ox4* and the gibberellin receptor genes *HvGID1L2* and *HvGID1L3*), *HvFT3* (also known as *PHOTOPERIOD 2*, *HvPPD-H2*), *EARLY FLOWERING 4-like4* (*HvELF4-like4*), and a homologue of *FLOWERING LOCUS T1* (*HvFT1-1*). More than 53% of the detected SNPs within these genes exhibited large differences in allele frequency among the different historical categories (≥20%; S4 Table). To investigate possible functions of all candidate genes, we performed gene ontology and pathway enrichment analyses, revealing that genes under selection during barley breeding were related to responses to oxidoreductase activity, peroxidase activity, and antioxidant activity (S16 Fig, S9 File).

We next predicted the variant effects of the 3,105 genetic variants located within the 69 candidate regions under selection in the 47 Australian barley cultivars within CatA, CatB, and CatC. Most genetic variants were detected in downstream (28%) or upstream gene regions (22%), while a large proportion of variants that fell within coding regions were missense variants (44%) (S17 Fig). Using a sorting intolerant from tolerant (SIFT) analysis, we identified 119 missense tolerated and deleterious mutations in 42 genes (S10 File). Of these variants, 29 were detected and annotated within 12 genes that exhibited large differences in allele frequency among the three historical categories (≥20%; S5 Table). These 12 genes—which based on the categories we conclude have been under selection during the past 120 years—were observed only on chromosome 7H. Hence, these genes are attractive candidates for further investigation, as they may hold the potential to enhance agronomic traits in Australian barleys through breeding.

### GWAS of agronomic traits

We field trialled and scored all 632 barley genotypes in the global barley panel for three key agronomic traits—flowering time (using days to Zadoks stage 49 [ZS49] as an equivalent for flowering time [22]), grain yield, and plant height—in sixteen independent field experiments at field sites located across the Western Australian wheatbelt region (Geraldton, Merredin, Katanning, Perth, and Esperance) conducted between 2015 and 2017. To evaluate the trait stability of the global barley panel across all locations and years, we calculated the coefficient of variation (CV) and heritability of each trait across all field trials (S11 and S12 Files). Z scores calculated per genotype for each field trial and trait revealed specific genotypes in the global barley panel with stable, consistent, and robust trait characteristics across the different field trials (S12 File). For example, the 2-row hulless and very early-maturing Canadian variety CDC Speedy was one of the earliest-flowering varieties, irrespective of location or year, whereas Spanish landrace 355 was consistently late-flowering and tall-growing across all environments and years.

We then performed a multi-environment and multi-year GWAS to test if genetic variation identified from the global barley panel is associated with the key agronomic traits flowering time, grain yield, and plan height. We used 33,486 filtered genetic markers with a MAF > 0.01 for GWASs based on two statistical models, generalized linear models (GLMs) and mixed linear models (MLMs). Manhattan plots and quantile-quantile (QQ) plots of the three traits are provided in S18–20 Figs. A graphical genotype map of selected significant genetic variants detected for all three traits is shown in Fig 5, and a similar map that includes P-values and marker *r^2^* values is shown in S21 Fig.

**Figure 5:**
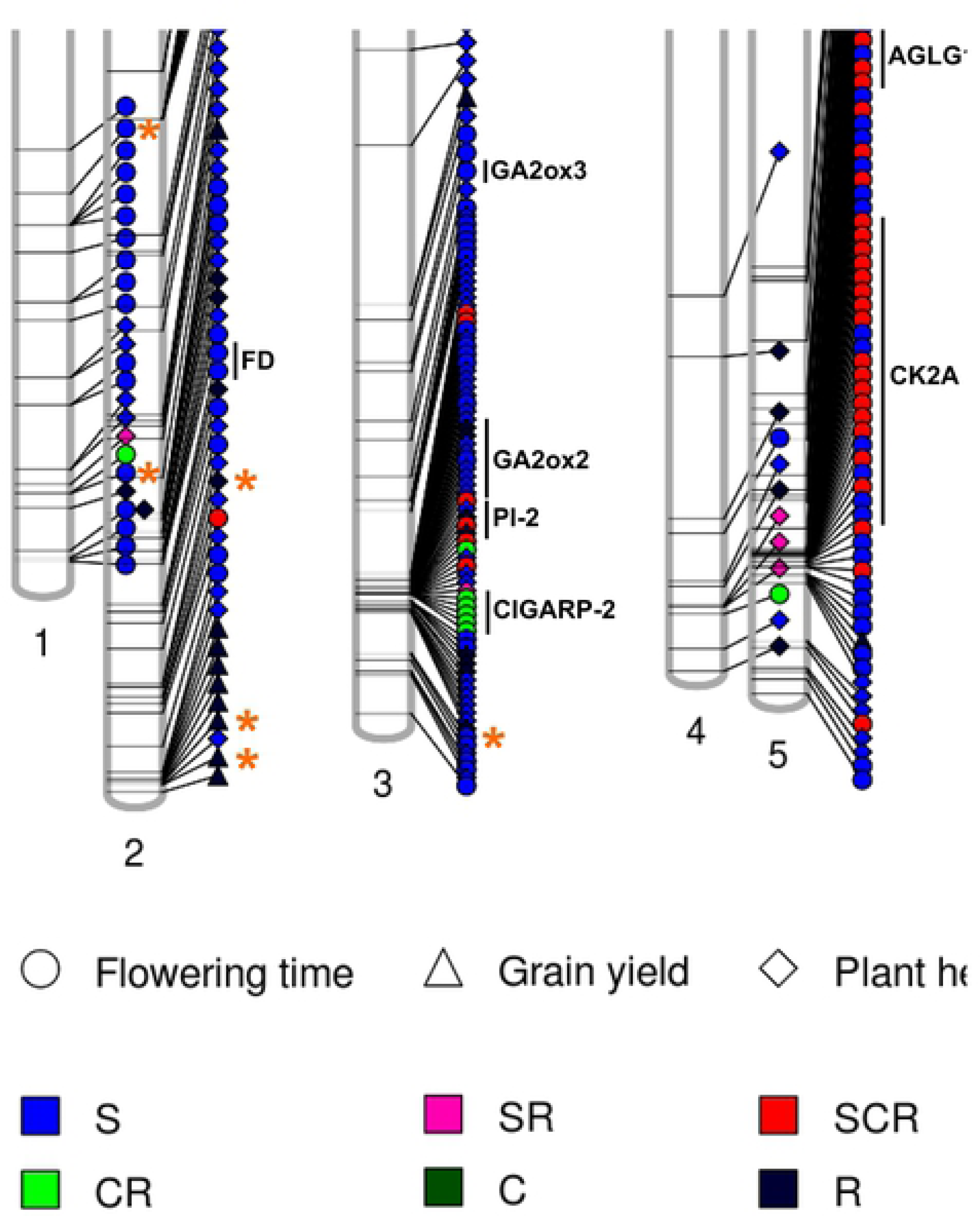
Graphical genotype map of selected genetic variants associated with three agronomic traits. Significant genetic variants detected via genome-wide association studies for flowering time (FT, measured as days to Zadoks stage 49 [ZS49]), grain yield (GY), and plant height (PH). Only stable, consistent, and/or robust markers are shown (Supplemental Files 13–15). Selected genetic variants (consistent and/or robust markers) that also fall within candidate regions for breeding selection are marked with orange asterisks. Plots drawn using the PhenoGram software tool.

First, we consider the genetic variation associated with flowering time. Across all field trials, we identified 1,132 significant unique marker-trait associations (MTAs) (false discovery rate [FDR] of P < 0.05) for flowering time located within 327 unique genes with functional annotations, each explaining up to 18.7% of the phenotypic variation (S13 File, S18 Figure). These regions include known phenology-related genes, such as *HvPPD-H1*, *PHYTOCHROME C* (*HvPhyC*), *PROTEIN KINASE 2A* (*HvCK2A*), *HvADA2, PHYTOCHROME-ASSOCIATED PROTEIN 2* (*HvPAP2*), and *VERNALIZATION H1* (*HvVRN-H1*) [4,10–12]. Furthermore, a total of 246 MTAs were considered to be ‘stable’, 76 MTAs were considered to be ‘consistent’, and 73 MTAs were considered to be ‘robust’ (S13 File). More than 30% of the significant MTAs were also detected in our previous study [10], in which a GWAS was performed using 4,600 SNPs (in comparison to the 33,486 SNPs used in the present study) obtained from target enrichment sequencing data for phenology genes combined with field trial data from 2015 and 2016. Here, novel and highly significant MTAs were located within the genes HORVU5Hr1G096560 (disease resistance protein), HORVU5Hr1G095040 (beta glucosidase C), and HORVU5Hr1G104240 (zinc finger A20 and AN1 domain-containing stress-associated protein 6) on chromosome 5H (S13 File).

Notably, we also detected novel associations with candidate phenology-related genes that were not included in the previous target-enrichment sequencing study [10] but that have annotations linking them to roles in flowering time regulation, including CCT and PRR motifs characteristic of key phenology genes such as *HvCO1*, *HvVRN-H2*, and *HvPPD-H1* [4]. These genes included HORVU1Hr1G011030 and HORVU5Hr1G125620 (both annotated as COP1-interacting protein-related), HORVU2Hr1G055130 and HORVU7Hr1G044380 (both annotated as CONSTANS, CO-like, and TOC1 [CCT] motif family protein, and HORVU3Hr1G092330 and HORVU6Hr1G008870 (both annotated as pentatricopeptide repeat-containing protein).

Next, we consider the genetic variation associated with grain yield. We identified 118 significant unique MTAs for grain yield, 30 of which were ‘robust’, with each MTA explaining up to 7.2% of the phenotypic variation (none were consistent or stable). We identified genetic variants within 30 functionally annotated genes (S14 File, S19 Fig). These regions included known phenology-related genes, such as *HvFT2* and *HvGA20ox2*. Like the results for flowering time, more than 30% of the significant MTAs for grain yield were also detected in our previous study [10]. Here, novel and highly significant MTAs were located within the genes HORVU2Hr1G125100 (peroxidase superfamily protein), HORVU7Hr1G002260 (disease resistance protein CC-NBS-LRR class family), and HORVU7Hr1G045290 (aluminum-activated malate transporter 9).

Finally, for plant height, we identified 1,279 significant unique MTAs within 395 functionally annotated genes, each explaining up to 8.9% of the phenotypic variation, including several gibberellin oxidase genes (*HvGA2ox4*, *HvGA20ox4*, and *HvGA2ox1*) and in particular the *sdw1/denso* gene (*HvGA20ox2*), a major determinant of plant height [23] (S15 File, S20 Fig). Furthermore, a total of 190 MTAs were considered to be ‘stable’, one MTA was considered to be ‘consistent’, and 61 MTAs were considered to be ‘robust’ (S14 File). Of these significant MTAs, approximately 8% were detected in our previous study [10]. Novel and highly significant MTAs were located within the genes HORVU3Hr1G021140 (gigantea protein GI), HORVU3Hr1G022170 (homeobox-leucine zipper protein ROC4), and HORVU3Hr1G089160 (AP2-like ethylene-responsive transcription factor). Our data also revealed relevant candidate genes for functional annotation that are known to have pleiotropic effects on several agronomic traits, as exemplified by the major flowering time and plant height associations detected on chromosome 5H, where *HvPhyC* was a major driver (S13 File), an association previously detected and discussed in detail [10].

Combining the GWAS with the results from the selective sweep analysis, we then compared genes containing MTAs for all three traits (flowering time, grain yield, and plant height) with genes located within the 69 candidate regions under selection that are related to breeding in Australian barley (S8 File). The results show that 23, 7, and 23 genes with significant MTAs for all three traits, respectively, are located within breeding-related genomic regions (S13–15 Files) including the known phenology genes *HvPPD-H1*, *AGAMOUS-LIKE GENE 1* (*HvAGLG1*), and *HvGA2ox3*. These results indicate that a subset of breeding loci are relevant for continued agronomic trait improvement and may have undergone additional selection to allow introduced European barleys to adapt to Australian growing conditions.

## Discussion

The conservation of genetic diversity for the future breeding of new crop varieties is particularly important in mitigating the adverse impacts of climate change on crop production. In this study, we assessed the contributions of both historical and recent breeding efforts towards local adaptation and crop improvement in a global barley panel of 632 genotypes. We used Australian barley as a model for the profound impact that breeding efforts had on developing a previously poorly adapted crop suitable to local environments.

Most Australian and international cultivars in the global barley panel were varieties that led the market when they were released. Thus, this study panel represents the long-term breeding progress for the world’s fourth-most, and Australia’s second-most widely grown crop at peak agronomic performance.

The current study employed ∼34,000 genetic markers—approximately 9 times more markers than used in previous studies of barley [10–12]—to perform diversity, selection footprint, and high-resolution GWAS analyses of agronomically relevant traits. The diversity analyses revealed that the most recently released Australian cultivars exhibited more than 12% higher nucleotide diversity than earlier-released cultivars. Thus, our results show that modern Australian barley cultivars are not genetically depauperate in comparison to historical varieties, which counters the common perception that intensive breeding leads to the erosion of adaptive genetic diversity in modern cultivars [1, 2]. This notion is supported by several recent reports on cereal crops such as wheat, which demonstrate that genetic diversity has not been reduced in European wheat cultivars over the past five decades of progress in breeding [24, 25]. Moreover, the faster LD decay and lower level of long-range LD of more recently released cultivars indicate that recent breeding efforts have increasingly integrated lines with more diverse genetic backgrounds into the pool of Australian germplasm. These observations also highlight the importance of breaking these large linkage blocks in future breeding efforts by increasing genetic diversity through new genetic crosses, such as unreleased breeding and research lines, landraces, and selected wild barley to eliminate the genetic hitchhiking of disadvantageous alleles within these LD blocks.

The selection footprint analyses for different Australian subpopulations detected substantial population differences (high F_ST_) and genomic regions with substantially lower levels of diversity in more recently released cultivars than in historical cultivars (CatB and CatC groups, high ROD) as possible candidate regions that were under selection during breeding in the recent past. We estimate that 2.3% of barley genes (i.e. 922 genes) fall into the selected category and thus have been affected by breeding selection in Australian barley. An excess of rare alleles relative to expectation for five phenology-related genes in historical Australian barley groups and seven phenology-related genes in geographic subpopulations are consistent with an increase in population size following a bottleneck or a selective sweep, and could indicate strong selection during post-domestication (breeding) diversification. Notably, a lack of rare alleles for the vernalisation response genes *VERNALISATION INSENSITIVE 3* (*HvVIN3*) and *HvZCCT-Hb* (the latter of which is a homologue of the *VERNALIZATION H2* (*VRN-H2*) locus) was detected for European and South American barley varieties, respectively. As the ancestor of domesticated barley is likely a winter-type wild barley [26], advantageous mutations in vernalisation response genes may have resulted in flowering time promotion in the absence of cold which facilitated the expansion of cultivable areas closer to the equator, pointing towards balancing selection for vernalisation requirements during past breeding efforts.

Next, we performed high-resolution GWAS analyses of agronomically relevant traits, and detected novel and highly significant MTAs for all three traits not detected in our previous study [10, 11]. For example, we detected novel and highly significant MTAs for plant height, located within the phenology genes HORVU3Hr1G021140 (gigantea protein GI), and HORVU3Hr1G022170 (homeobox-leucine zipper protein ROC4). ROC4 regulates the transcript levels of *GRAIN NUMBER, PLANT HEIGHT, AND HEADING DATE7* (*Ghd7*) and causes long day-dependent early flowering in rice [27], whereas *GI* is a circadian clock-controlled gene responsible for fine-tuning plant developmental processes in response to photoperiod [28]. For grain yield, novel and highly significant MTAs were located within the genes HORVU2Hr1G125100 (peroxidase superfamily protein), HORVU7Hr1G002260 (disease resistance protein CC-NBS-LRR class family), and HORVU7Hr1G045290 (aluminum-activated malate transporter 9), which all play functional roles in pathogen and abiotic stress resistance [29, 30]. Interestingly, a gene encoding a CC-NBS-LRR-class disease resistance protein was recently reported to be associated with grain yield in chickpea [31].

Finally, the diversity, selection footprint, and GWAS analyses performed in the present study demonstrate that several key loci, including major phenology genes such as *HvPPD-H1*, harbour coincident signals, supporting the view that a subset of breeding loci is relevant to the continued improvement of agronomic traits and have undergone additional selection in the adaptation of introduced European barleys to Australian growing conditions. It will be interesting to follow-up these results with detailed genetic analyses on individual genes to characterize their functions in more detail.

In summary, our combined variant dataset and germplasm collection provide a rich source of genetic information that can be applied to understanding and improving diverse traits, such as environmental adaptation and enhanced yield, and could accelerate genetic gains in future barley breeding.

## Materials and Methods

### Ethics Statement

The research for this project does not require ethics approval in Australia. All data are available in the supplementary documents and public database.

### Plant material

The barley panel consisted of 632 genotypes, including 250 cultivars and 382 breeding and research accessions from 37 countries throughout Europe, Asia, North and South America, Africa, and Australia, and were selected from over 4,000 accessions preserved at the Western Barley Genetics Alliance at Murdoch University (Perth, Australia) to represent barley genotypes from major global barley breeding programmes. For a detailed description of all lines and varieties used in this study, see S1 File. This panel spanned the entire spectrum of cultivated barley, consisting of two-(92%) and six-row (8%) genotypes, and of winter (7%), spring (92%), and facultative (1%) growth habits. The selected germplasm also included 47 Australian cultivars released since 1903 as well as 180 Australian breeding and research lines. All cultivars of the global barley panel are highly productive and genetically uniform commercial varieties developed by professional plant breeders, whereas the all breeding and research lines include germplasm collections for potential use in developing future cultivars.

### Field experiments and phenotypic data

A total of 16 field experiments were conducted in 2015, 2016, and 2017 across a variety of environments in Western Australia (South Perth, Geraldton, Katanning, two sites at Merredin and Esperance, respectively. The number of cultivars of the global barley panel tested at each field site are provided in S11 File. In Western Australia, Geraldton, South Perth, and Esperance are located along the coast of the Southern Ocean and all receive high annual rainfall but have very different daily maximum temperatures (the Geraldton site being the warmest, and the Esperance site being the coolest). The distance between Geraldton and Esperance is over 1,100 km. The Merredin site is located inland and receives little rainfall, while the Katanning site receives a medium amount of rainfall. The experimental design for field trial sites was performed as previously described [10]. Briefly, all regional field trials (partially replicated design) were planted in a randomized complete block design using plots of 1 by 3 m^2^ laid out in a row-column format. Field trials in South Perth were conducted using a hill plot technique with a 40-cm distance within and between rows due to space limitations. Seven control varieties were used for spatial adjustment of the experimental data.

Measurements were taken at each plot of each field experiment in the study to determine flowering time (days to ZS49), plant height, and grain yield as previously described [10]. Briefly, plant maturity was recorded as the number of days from sowing to 50% awn emergence above the flag leaf (ZS49) [32], as a proxy for flowering time [22]. Plant height was determined by estimating the average height from the base to the tip of the head of all plants in each plot. Grain yield (kg ha^−1^) was determined by destructively harvesting all plant material from each plot to separate the grain and determining grain mass. Grain yield data collected in the 2015, 2016, and 2017 field experiments, as well as plant height and plant maturity data for the 2017 field trials, were analysed using linear mixed models (LMMs) in ASReml-R (https://www.vsni.co.uk/software/asreml-r/) to determine best linear unbiased predictions (BLUPs) or best linear unbiased estimations (BLUEs) for each trait for further analysis. Local best practices for fertilization and disease control were adopted for each trial site.

### Evaluation of agronomic trait stability across field sites and genotypes

To evaluate the yield stability of global barley panel across all location and year scenarios in the main trials, we calculated the CV for flowering time, grain yield, and plant height across all location-by-year combinations, along with Z scores according to Equation 1:

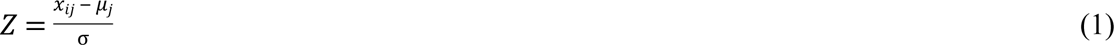

where x_ij_ is the trait value of variety i for year-by-location combination j, μ_j_ is the mean of the trait value for all plants of variety i in j, and σ is the standard deviation of the population mean.

Z scores were used to determine above-(positive Z score) and below-average (negative Z score)-yielding cultivars, as well as cultivars that flowered earlier (negative Z score) or later (positive Z score) or were shorter (negative Z score) or taller (positive Z score) than average for all year-by-location combinations. The critical Z score values for a 95% confidence level were −1.96 and +1.96 standard deviations, equal to a P-value of 0.05. Genotype trait characteristics (e.g. early flowering, high yielding, or short stature) were defined as ‘robust’ if they were consistently below or above the population mean in one location, ‘stable’ if they were significant (less than −1.96 or greater than +1.96 standard deviation) in more than one location, and ‘consistent’ if they were significant (less than −1.96 or greater than +1.96 standard deviation) across at least two years at one or more locations.

### DNA extraction

Genomic DNA was extracted from the leaves of a single barley plant per variety using the cetyl-trimethyl-ammonium bromide (CTAB) method as previously described [10, 11]. DNA quality was assessed on 1% agarose gels and quantified using a NanoDrop spectrophotometer (Thermo Scientific NanoDrop Products, Wilmington, Delaware USA).

### Sequencing, sequence alignment, genotype calling, variant discovery, and variant prediction

As the barley genome is quite large (∼5.1 Gb), and the genome consists of >80% mobile and repeated elements, whole-genome re-sequencing is a cost-intensive approach to comprehensively catalogue genetic diversity. To circumvent this limitation, we used a combination of three sequencing methods (target-enrichment sequencing, low-coverage WGS, and DArTseq) to capture variation in and around the gene-containing regions of the 632 barley genotypes.

### Target-enrichment sequencing

To assess the genetic diversity of phenology and phenology-related genes in the global collection of barley landraces and cultivars, we designed a custom target-enrichment sequencing assay for loci implicated in the flowering pathway in barley and related plant species, as previously described [10, 11]. In short, the target-enrichment sequencing of genomic DNA regions was performed by solution-based hybrid capture using a synthetic library consisting of 13,588 RNA probes (MYbaits, MYcroarray®, Ann Arbour, MI, USA) following the manufacturer’s protocol (v.2.3.1). Post-capture DNA libraries were combined into 10 pools of approximately 96 samples each and sequenced on three lanes on an Illumina HiSeq 3000 (Illumina Inc., San Diego, CA, USA) to generate approximately 0.5 million 2x150-bp paired-end reads per sample. Genome sequencing was conducted at AgriBio (Centre for AgriBioscience, Bundoora, VIC, Australia). Sequence files were post-run filtered and aligned to the latest release of the barley reference genome assembly [7] (IBSC v2) using Nuclear software v.3.6.16 (GYDLE Inc., Montreal, Canada). SNP variant discovery and genotype calling were performed using custom Perl scripts to produce a variant call format (VCF) v.4.2 genotype file based on the alignment files as previously described [10, 11]. Only SNPs with <10% missing values and a MAF >1% (4,260 SNPs) were used for subsequent analyses.

### Low-coverage whole genome sequencing

Each sample for low-coverage (1x) WGS consisted of a pool of 20 individual barley pre-capture DNA libraries from the target-enrichment sequencing experiment, dissolved in 10 mM Tris HCl (pH of 8.0). Thirty microliters of the pooled remainders of pre-capture DNA libraries were subjected to low-coverage WGS by the Beijing Genome Institute (BGI, Hong Kong) on an Illumina HiSeq 4000 to generate approximately 50 million 2x150-bp paired-end reads per sample. The latest release of the barley reference genome assembly (IBSC v2) was used as a reference to map the clean reads with the alignment algorithm BWA-MEM [33] using default parameters. Duplicates were marked and removed using Picard v.1.129 (http://broadinstitute.github.io/picard/). Only reads with unique mapping positions in the reference genome were retained and used to detect genomic variations (SNPs and InDels).

InDels and SNPs were detected by running three rounds of SAMtools v.1.7 plus BCFtools v.1.7 and the Genome Analysis ToolKit [34] (GATK v.3.8) variant-calling pipeline. Briefly, the first round was performed with SAMtools plus BCFtools, with filtering based on both mapping quality and variant calling quality. The result of the first round was used as a guide for realignment around potential InDels and variant calling for the second round of GATK. The common variants detected by both the SAMtools plus BCFtools pipeline and the GATK pipeline were used to guide variant calling in the third round using GATK.

### Genotyping-by-sequencing by DArTseq

In addition, DArTseq GBS was performed using the DArTseq platform (DArT PL, Canberra, NSW, Australia) according to the manufacturer’s protocol (https://www.diversityarrays.com/). Briefly, 100 μl of 50 ng μL^−1^ DNA was sent to DArT PL, and GBS was performed using complexity reduction followed by sequencing on a HiSeq Illumina platform as previously described [35]. DArTseq marker sequences were aligned against the Morex barley genome assembly [7] IBSC v.2. The genetic position of each marker was determined based on the Morex physical reference assembly. Filtered DArTseq GBS (<10% missing values and minor allele frequency [MAF] >1%) yielded 14,032 SNPs across all 632 samples.

Stringent filtering steps were adopted to obtain clean data as previously described [10, 11]. All genotype data were combined, filtered based on duplicates and MAF >1%, and imputed using BEAGLE v.4.1 [36] to yield a final number of 33,486 filtered genetic markers (32,645 SNPs and 841 InDels) with a MAF > 1%.

### Population structure and genotype data analyses

As previously described [10], the PIC was calculated for each of the 33,486 filtered genetic markers according to Equation 2:

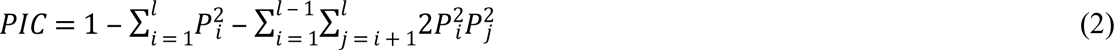

where Pi and Pj are the population frequencies of the i^th^ and j^th^ alleles, respectively.

The ADMIXTURE v.1.3.0 model-based clustering algorithm [19] was used to investigate the subpopulation structure of the global barley panel. Prior to subpopulation structure analysis in ADMIXTURE, the genotype dataset was LD pruned using Plink v1.9 [37] with a window size of 50 kb, step size of 5, and pairwise *r^2^* threshold of 0.5, yielding 18,869 genetic variants. A preliminary analysis was performed using 100 replicate runs by inputting successive values of K from 1 to 18, as previously described [10]. A 10-fold cross-validation procedure was performed with 100 different fixed initial seeds in multi-threaded mode for each K-value. The most likely K-value was determined using ADMIXTURE cross-validation error values.

CLUMPP [38] v.1.1.2 software was used to obtain the optimal alignments of 100 replicates for each K-value. Membership proportions of each genotyped individual were averaged across runs according to the permutation with the greatest symmetric similarity coefficient as described previously [10]. The output from CLUMPP for the optimal K-value was used to make plots using the cluster visualization package Pophelper v.2.2.3 [39] implemented in R v.3.5.1 (http://www.R-project.org/).

To summarize the genetic structure and variation present in the barley germplasm, PCA was also conducted using all 33,486 filtered genetic markers in TASSEL [40] v.5.2.39. The first three PCs were plotted against each other using the ‘scatter plot’ function in Microsoft Excel 2016. NJ trees were constructed using the Java application Archaeopteryx v.0.9909 [41] based on genetic distances calculated in TASSEL v.5.2.39. The sub-structures in the collection inferred using different methodologies were compared, and the final K-value was ascertained using ADMIXTURE [19].

### Linkage disequilibrium

Genome-wide LD analysis was performed for the global barley panel and subgroups using all 33,486 filtered genetic markers using Plink v.1.9 [37]. LD was estimated by using squared allele frequency correlations (*r^2^*) between the intra-chromosomal pairs of loci [42]. The loci were considered to be in significant LD when P < 0.001. To investigate the extent of and average LD decay in the panel, significant inter- and intra-chromosomal *r^2^* values within each 100-kb bin were plotted against the physical distance (kb) between markers. Curves were fitted by a second-degree LOESS function using R v.3.5.1 (http://www.R-project.org/).

### Diversity parameter estimation and detection of selective sweeps

To detect genomic areas with selective sweeps driven by artificial (breeding) selection, we calculated F_ST_, π, and ROD using VCFtools v.0.1.14 [43] and a window size of 10 Mb. F_ST_ estimates for pairs of subpopulations were calculated as previously described [44].

Subpopulation-specific estimates of Tajima’s D were calculated using VCFtools v.0.1.14 [43] to compare the average number of pairwise differences and the number of segregating sites between samples within each of our geographic and historical subpopulations in Australia.

The ROD index was calculated for each 10-Mb window based on the ratio of diversity between Australian subpopulation CatB to that of CatA according to Equation 3:

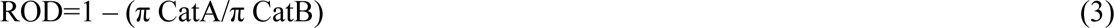

and between Australian subpopulation CatC to that of CatA according to Equation 4:

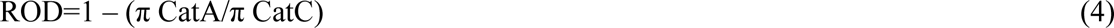

where the nucleotide diversity statistic π is the average number of nucleotide differences between any two DNA sequences. In addition, whole-genome screening of selected regions was performed using XP-CLR, a likelihood method for detecting selective sweeps that is based on the multilocus allele frequency differentiation between two populations [21]. XP-CLR tests were run with a window size and step size of 1 Mb, with CatA set as the reference and compared to CatB and CatC for each chromosome. Invariant or singleton SNPs were excluded, leaving on average ∼45% of available variants for the analysis.

A total of 69 regions, which were in the highest 95^th^ (F_ST_, ROD) or 99^th^ (XP-CLR) percentile of all regions identified, were considered to be under selection. Within the identified 69 regions under selection, 3,105 genetic variants were located within 922 genes. The 3,105 genetic variants were used for VEP using the Ensembl Variant Effect Predictor toolset (Ensembl Variant Effect Predictor web interface, http://www.ensembl.org/vep). VEP was performed to determine the effect of the genetic variants on genes, transcripts, and protein sequence, as well as regulatory regions. SIFT was estimated to predict the effects of amino acid substitutions on protein function based on sequence homology and the physical properties of amino acids. The results were filtered for missense only with SIFT scores (SIFT score <0.05) from tolerant to deleterious. Regions of genetic differentiation between subpopulations and genes within these regions were identified based on F_ST_, π, ROD, and XP-CLR values of markers plotted linearly along each chromosome according to physical position. All visualizations were performed using the R packages yarrr and ggplot2.

### Gene ontology and pathway enrichment analysis

Gene ontology and pathway enrichment analysis of the 922 candidate genes under selection was performed as previously described [45]. Briefly, singular enrichment analysis (SEA) was performed using AgriGO v.2.0 [46] with the following parameter settings: Fisher’s test, 0.05 significance level, 5 minimum mapping entries, and complete gene ontology type.

### Association analysis

GWASs were performed using TASSEL v.5.2.39 [40] and a total of 33,486 filtered genetic variants with <10% missing values and a MAF >1%. Different statistical models were used to calculate P-values for putative MTAs as follows, which included population structure to avoid spurious associations. For the 2015 and 2016 data, a compressed MLM with a population structure (Q) matrix (PCs) and kinship (K) matrix (matrix of genetic similarities based on simple matching coefficients) was used to correct for population structure as previously described [10]. According to the QQ plot, the MLM that incorporated Q and K was suitable for these datasets. Data from the 2015 and 2016 field trials were used in a previously published GWAS using only target-enrichment sequencing data (4,260 SNPs) [10]. For the 2017 data, GLMs with PCs as a correction for population structure were tested for all associations, which, according to the QQ plots, were suitable for this study. For all MTAs, multiple testing using Storey’s q-value method [47] was performed to control for false discoveries and to assess statistical significance. As part of the q-value method, the smoother method, an extension of FDR correction, was employed. Lambda was set to 0, which estimates πi(0) =1, produces a list of significant tests equivalent to that obtained with the Benjamini and Hochberg [48] procedure, and is considered to be a conservative case of Storey’s q-value methodology. Only markers with a qFDR of <0.05 were considered to be significant. Manhattan plots were drawn with qman [49] v.0.1.4.

Broad-sense heritability (*H^2^*) was calculated using the following equation by treating genotype and environment as random effects, applying an MLM according to Equation 5:

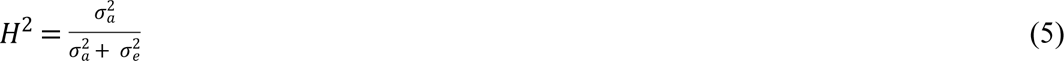

where σ^2^ and σ^2^ represent the variance derived from genotypic and environmental effects, respectively.

MTAs were defined as ‘robust’ if they explained more than 5% of the phenotypic variation, ‘stable’ if they were identified in more than one location, and ‘consistent’ if they were identified in more than one year. Phenogram [50] was used to produce the graphical genotype map in Fig 7.

## Acknowledgements

We thank Ms Lee-Anne McFawn and Mr David Farleigh from DPIRD (South Perth, WA) for providing technical assistance in the field trials.

## Supporting Information captions

**S1 Fig. Barley genotype panel.** The barley diversity panel consists of 632 genotypes sourced worldwide a), with genotypes separated by b) continents of origin (Africa, North America, South America, Asia, Europe and Australia), c) row type, and d) growth habit. Insert figure within b): Historical categories are presented for Australian cultivars only (Cat A: historic cultivars, released prior to 1999; Cat B: modern cultivars, released between 1999 and 2005; Cat C: recent cultivars, released between 2006 and 2019; and Cat D: breeding and research lines).

**S2 Fig. Distribution of genetic variant density on seven chromosomes.** Number of genetic variants (y-axis) for the seven chromosomal groups over 10 Mb sliding windows (x-axis).

**S3 Fig. Graphical genotype map of genetic variants detected using different genotyping technologies.** Genetic markers detected via a) low-coverage whole genome sequencing, b) Genotyping-by-sequencing (GBS) by Diversity Arrays Technology sequencing (DArT-Seq), c) target capture sequencing, and d) all genetic markers combined (a- c). Plots drawn using the PhenoGram online software tool. Cat C: recent cultivars, released between 2006 and 2019; and Cat D: breeding and research lines).

**S4 Fig. Exploration of the optimal number of genetic subpopulations (K) using Δ cross-validation error and standard error values in the barley germplasm collection.** A solid line denotes the choice of K=12 which represents the most likely number of subpopulations within the barley germplasm collection.

**S5 Fig. Plot of ancestry estimates inferred by ADMIXTURE for 632 worldwide barley accessions.** Each colour represents a population, and the colour of individual haplotypes represents their proportional membership in the different populations. Membership coefficients for each population were merged across 100 replicate runs using the CLUMPP programme. The number of clusters (K) present in the entire population of 632 accessions was judged to be K = 12 based on the CV error. Shown are clusters 2, 4, 6, 8, 10, and 12.

**S6 Fig. Neighbour-joining trees of 47 selected Australian barley cultivars.** Each colour represents a different historic group as per legend. The tree was constructed from simple matching distance of 33,486 common genetic variants in the selected barley cultivars.

**S7 Fig. Principal component analysis (PCA) of the first two components of 632 barley varieties.** a) PCA based on geographic region: The seven divergence groups are coloured respectively; b) PCA based on growth habit: The three divergence groups are coloured respectively; c) PCA based on row type: The two divergence groups are coloured respectively. PC1 and PC2 together explain about 54% of the total variation, and partitioned the population into distinct clusters.

**S8 Fig. The extent of Linkage Disequilibrium (LD) across the seven chromosomes in a worldwide collection of domesticated barley varieties.** Values are intra-chromosomal LD r^2^ values for all intra-chromosomal pairs of genetic variants binned by distance. Curves were fitted by second-degree LOESS curve.

**S9 Fig. Chromosomal distribution of nucleotide diversity (π) between different historic groups of domesticated Australian barley genotypes.** Statistics based on 10 Mb windows. A legend is provided at the top of the figure. CatA: Cultivars released between 1903 and 1998; CatB: Cultivars released between 1999 and 2005; CatC: Cultivars released between 2006 and 2019.

**S10 Fig. Chromosomal distribution of nucleotide diversity (π) between different geographic groups of the barley diversity panel.** Statistics based on 10 Mb windows. A legend is provided at the top of the figure.

**S11 Fig. Chromosomal distribution of Tajima’s D values between different historic groups of domesticated Australian barley genotypes.** Statistics based on 10 Mb windows. Filled circles show values above the 99th percentile and are colour-coded according to the different historic groups. A legend is provided at the top of the figure. CatA: Cultivars released between 1903 and 1998; CatB: Cultivars released between 1999 and 2005; CatC: Cultivars released between 2006 and 2019. CatD: unreleased breeding and research lines. legend is provided at the top of the figure.

**S12 Fig. Chromosomal distribution of Tajima’s D values between different geographic groups of the barley diversity panel. Statistics based on 10 Mb windows.** Filled circles show values above the 99th percentile and are colour-coded according to the different geographic groups. A legend is provided at the top of the figure.

**S13 Fig. Chromosomal distribution of Fixation Index (FST) values and XP-CLR scores between different historic groups of domesticated Australian barley genotypes.** Highlighted regions (triangles coloured as per legend) are based on the 99th percentile (XP-CLR). Statistics based on 10 Mb windows. A legend is provided at the top of the figure. CatA: Cultivars released between 1903 and 1998; CatB: Cultivars released between 1999 and 2005; CatC: Cultivars released between 2006 and 2019.

**S14 Fig. Chromosomal distribution of Reduction of Diversity (ROD) values between CatA and CatB historic groups of domesticated Australian barley genotypes.** Statistics based on 10 Mb windows. A legend is provided at the top of the figure. CatA: Cultivars released between 1903 and 1998; CatB: Cultivars released between 1999 and 2005. 2005; CatC: Cultivars released between 2006 and 2019.

**S15 Fig. Chromosomal distribution of Reduction of Diversity (ROD) values between CatA and CatC historic groups of domesticated Australian barley genotypes.** Statistics based on 10 Mb windows. A legend is provided at the top of the figure. CatA: Cultivars released between 1903 and 1998; CatC: Cultivars released between 2006 and 2019.

**S16 Fig. Significantly enriched GO terms related to molecular function in genes within candidate regions under selection.** Gene Ontology (GO) and Pathway enrichment analysis performed with AgriGO v.2.0 using Fisher test, 0.05 significance level, 5 minimum mapping entries and Complete GO gene ontology type. Full datasets are available in S9 File.

**S17 Fig. Consequences of genetic polymorphisms identified in candidate regions under selection and categorized by Ensembl Variant Effect Predictor.** A total of 3,105 genetic variants are categorized and percentages of potential consequences are provided in a) for all consequences, and b) for consequences in coding regions only. See Ensembl Variant documentation for explanation of consequence categories (http://www.ensembl.org/info/genome/variation/predicted_data.html#consequences).

**S18 Fig. Manhattan and QQ plots of flowering time for all field trials with significant marker-trait associations.** Days to ZS49 were used as an equivalent to flowering time (FT). Left panel: Manhattan plots, right panel: Quantile–quantile (QQ) plots. GWAS results are presented by negative log_10_ of unadjusted p-values against position on each of the seven chromosomes. Horizontal dashed lines indicate the genome-wide significant threshold selected by local false discovery rate and a q-value cut-off at 0.05 (blue) and 0.01 (red).

**S19 Fig. Manhattan and QQ plots of grain yield for all field trials with significant marker-trait associations.** Other details as per legend to S18 Fig.

**S20 Fig. Manhattan and QQ plots of plant height for all field trials with significant marker-trait associations.** Other details as per legend to S18 Fig.

**S21 Fig. Graphical genotype map of selected genetic variants including significant p-values and marker r2 values associated with three agronomic traits.** Significant genetic variants detected via GWAS for a) flowering time (measured as Days to ZS49), b) grain yield, and c) plant height. Only stable, consistent and/or robust markers are shown (S15 File).

**S1 Table. Barley genetic variants.** Sequencing of the 632 genotypes delivered 33,486 filtered genetic markers detailed in the current table with the number of variants (SNPs and InDels) from low-coverage whole genome sequencing (LC), target capture sequencing (TC), and DArTseq (DArT), the number of genic and non-genic variants and associated ratio (genic/non-genic) with the number of targeted genes per chromosomes. ‘Genic variants’ columns count include those associated with a high confidence annotation7. ‘Genes’ column includes genes with high-confidence annotation. ‘Non-genic’ variants include both ‘Upstream’ and ‘Downstream’ variants as well as variants without associated high- confidence annotation.

**S2 Table. Nucleotide diversity index (π) and Tajima’s D summary statistics of the genetic variation observed within different subpopulations in the barley diversity panel.** CatA: Cultivars released between 1903 and 1998; CatB: Cultivars released between 1999 and 2005; CatC: Cultivars released between 2006 and 2019.

**S3 Table. FST summary statistics in the genetic variation observed between different subpopulations in the barley diversity panel.** CatA: Cultivars released between 1903 and 1998; CatB: Cultivars released between 1999 and 2005; CatC: Cultivars released between 2006 and 2019.

**S4 Table. Allele frequency of missense genetic variants in phenology-gene related candidate genomic regions that underwent selection during breeding of in Australian barley cultivars.** Most phenology-gene related candidate genomic regions were part of previously published targeted phenology gene re-sequencing performed on the same barley cultivars10,11. Shown are only variants within phenology-gene related gene regions (including 500bp flanking regions) and with >20% allele frequency (freq.) difference between CatA, CatB, and CatC barley cultivars. TF: transcription factor. CatA: Cultivars released between 1903 and 1998; CatB: Cultivars released between 1999 and 2005; CatC: Cultivars released between 2006 and 2019.

**S5 Table. Allele frequency of missense genetic variants in candidate genomic regions that underwent selection during breeding of in Australian barley cultivars.** Shown are only variants with >20% allele frequency (freq.) difference of all missense variants between CatA, CatB, and CatC barley cultivars. All candidate genomic regions are located on chromosome 7H. CatA: Cultivars released between 1903 and 1998; CatB: Cultivars released between 1999 and 2005; CatC: Cultivars released between 2006 and 2019.

**S1 File. List of barley genotypes.** The barley diversity panel consists of 632 genotypes sourced worldwide.

**S2 File. Genome-wide Linkage Disequilibrium (LD) decay for all barley genotypes.** Values are mean and median LD r^2^ values for all pairs of genetic variants binned by distance (100kb).

**S3 File. Genome-wide Linkage Disequilibrium (LD) decay for Australian barley cultivars released between 1903 and 1998 (CatA).** Values are mean and median LD r^2^ values for all pairs of genetic variants binned by distance (100kb).

**S4 File. Genome-wide Linkage Disequilibrium (LD) decay for Australian barley cultivars released between 1999 and 2005 (CatB).** Values are mean and median LD r2 values for all pairs of genetic variants binned by distance (100kb).

**S5 File. Genome-wide Linkage Disequilibrium (LD) decay for Australian barley cultivars released between 2006 and 2019 (CatC).** Values are mean and median LD r^2^ values for all pairs of genetic variants binned by distance (100kb).

**S6 File. Genome-wide Linkage Disequilibrium (LD) decay for Australian barley breeding and research lines (CatD).** Values are mean and median LD r^2^ values for all pairs of genetic variants binned by distance (100kb).

**S7 File. Selected candidate regions from comparisons of Australian barley cultivars released between 1903 and 2019.** There are 69 selected regions from comparing CatA withCatB and CatC, respectively, and 924 candidate genes falling within the regions. CatA: Cultivars released between 1903 and 1998; CatB: Cultivars released between 1999 and 2005; CatC: Cultivars released between 2006 and 2019. Region ID: The identifier of a selected region; Region Coordinate: The range of the selected region; F_ST_ (CatA-CatB): Fixation Index (F_ST_) values among CatA and CatB barley historic groups in the candidate region. Highlighted regions are above the 95^th^ percentile. F_ST_ (CatA-CatC): Fixation Index (FST) values among CatA and CatC barley historic groups in the candidate region. Highlighted regions are above the 95^th^ percentile. ROD (CatA-CatB): Reduction Of Diversity (ROD) values among CatA and CatB barley historic groups in the candidate region. Highlighted regions are above the 95^th^ percentile. ROD (CatA-CatC): Reduction Of Diversity (ROD) values among CatA and CatC barley historic groups in the candidate region. Highlighted regions are above the 95^th^ percentile. XP-CLR (CatA-CatB): Cross-Population Composite Likelihood Ratio Test (XP-CLR) values among CatA and CatB barley historic groups in the candidate region. Highlighted regions are above the 99^th^ percentile. XP-CLR (CatA-CatC): Cross-Population Composite Likelihood Ratio Test (XP-CLR) values among CatA and CatC barley historic groups in the candidate region. Highlighted regions are above the 99^th^ percentile.

**S8 File. Candidate genes within selected candidate regions from comparisons of Australian barley cultivars released between 1903 and 2019.** There are 69 selected regions from comparing CatA with CatB and CatC, respectively, and 924 candidate genes falling within the regions. Of these, 890 have gene-stable ID’s and additional information is provided in this dataset. CatA: Cultivars released between 1903 and 1998; CatB: Cultivars released between 1999 and 2005; CatC: Cultivars released between 2006 and 2019. Gene ID: The candidate gene locus from barley annotation of IBSC v2 [4]; Gene Type: A gene classification; Gene Coordinate: The range of the gene from annotation; Functional Annotation: Annotation from IBSC v2, Gene name (TC): Abbreviated name given for genes selected and sequenced in the targeted re-sequencing (TC) phenology gene project [10,11].

**S9 File. Gene Ontology and pathway enrichment analysis for candidate genes under selection in Australian barleys.** Singular Enrichment Analysis (SEA) was performed using AgriGO v.2.0 using Fisher test, 0.05 significance level, 5 minimum mapping entries and Complete GO gene ontology type.

**S10 File. Variant Effect Predictor analysis for candidate genes under selection.** Variant Effect Predition (VEP) was performed using ensembl (McLaren et al., 2016) for candidate genes potentially under selection. Shown are results for missense only with Sorting Intolerant From Tolerated (SIFT) scores.

**S11 File. Descriptive statistics for all investigated traits in the field trials.** Minimal (Min) and Maximal (Max) values are shown next to the arithmetic mean (Mean), Standard deviation (SD) and coefficient of variation (CV). H^2^: Broad-sense heritability. N: Number of lines scored successfully in the field trial. *Plant development measured on 2 Oct as an estimation to ZS49. Esperance (1): Esperance field trial location 1. Esperance (2): Esperance field trial location 2 (EDRS). Merredin (1): non-irrigated. Merredin (2): irrigated. Perth (1): Perth, time of sowing 1 (early). Perth (2): Perth, time of sowing 2 (mid). Perth (3): Perth, time of sowing 3 (late).

**S12 File. Z scores for all barley varieties and investigated traits in the field trials.** The critical Z score values for a 95% confidence level were -1.96 and +1.96 standard deviations, equal to a P-value of 0.05. Genotype trait characteristics were defined as ‘robust’ if they were consistently below or above the population mean across one location (same value direction), ‘stable’ if they were significant (less than -1.96 or more than +1.96 standard deviations) for more than one location, and ‘consistent’ if they were significant (less than -1.96 or more than +1.96 standard deviations) across at least two years at one location or more. *Plant development measured on 2 Oct as an estimation to ZS49. Esperance (1): Esperance field trial location 1. Esperance (2): Esperance field trial location 2 (EDRS). Merredin (1): non- irrigated. Merredin (2): irrigated. Perth (1): Perth, time of sowing 1 (early). Perth (2): Perth, time of sowing 2 (mid). Perth (3): Perth, time of sowing 3 (late).

**S13 File. Significant marker-trait associations for flowering time (FT, Days to ZS49) identified via genome-wide association mapping for all studied agronomic traits.** MAF: Minor allele frequency. R2: Contribution to phenotypic variation. P-value: Adjusted p-value after multiple comparisons false discovery rate testing using Storey’s qvalue (Storey, 2002). Esperance (1): Esperance field trial location 1. Esperance (2): Esperance field trial location 2 (EDRS). Merredin (1): non-irrigated. Merredin (2): irrigated. Perth (1): Perth, time of sowing 1 (early). Perth (2): Perth, time of sowing 2 (mid). Perth (3): Perth, time of sowing 3 (late). Stable: Marker-trait association (MTA) detected at more than one location. Consistent: MTA detected for more than one year at the same location. Robust: More than 5% phenotypic variation explained.

**S14 File. Significant marker-trait associations for grain yield identified via genome-wide association mapping for all studied agronomic traits.** MAF: Minor allele frequency. R2: Contribution to phenotypic variation. P-value: Adjusted p-value after multiple comparisons false discovery rate testing using Storey’s qvalue (Storey, 2002). Esperance (1): Esperance field trial location 1. Merredin (1): non-irrigated. Stable: Marker-trait association (MTA) detected at more than one location. Consistent: MTA detected for more than one year at the same location. Robust: More than 5% phenotypic variation explained.

**S15 File. Significant marker-trait associations for plant height identified via genome- wide association mapping for all studied agronomic traits.** MAF: Minor allele frequency. R2: Contribution to phenotypic variation. P-value: Adjusted p-value after multiple comparisons false discovery rate testing using Storey’s qvalue (Storey, 2002). Esperance (1): Esperance field trial location 1. Merredin (1): non-irrigated. Stable: Marker-trait association (MTA) detected at more than one location. Consistent: MTA detected for more than one year at the same location. Robust: More than 5% phenotypic variation explained

